# Modeling of histone modifications reveals formation mechanism and function of bivalent chromatin

**DOI:** 10.1101/2021.02.03.429504

**Authors:** Wei Zhao, Lingxia Qiao, Shiyu Yan, Qing Nie, Lei Zhang

## Abstract

Bivalent chromatin is characterized by occupation of both activating histone modifications and repressive histone modifications. While bivalent chromatin is known to link with many biological processes, the mechanisms responsible for its multiple functions remain unclear. Here, we develop a mathematical model that involves antagonistic histone modifications H3K4me3 and H3K27me3 to capture the key features of bivalent chromatin. Three necessary conditions for the emergence of bivalent chromatin are identified, including advantageous methylating activity over demethylating activity, frequent noise conversions of modifications, and sufficient nonlinearity. The first condition is further confirmed by analyzing the experimental data from a recent study. Investigation of the composition of bivalent chromatin reveals that bivalent nucleosomes carrying both H3K4me3 and H3K27me3 account for no more than half of nucleosomes at the bivalent chromatin domain. We identify that bivalent chromatin not only allows transitions to multiple states but also serves as a stepping stone to facilitate a step-wise transition between repressive chromatin state and activating chromatin state, and thus elucidate crucial roles of bivalent chromatin in mediating phenotypical plasticity during cell fate determination.

## Introduction

A variety of covalent modifications on histones, such as methylation and acetylation, play important roles in governing the chromatin state. Such diversity of histone modifications confers additional complexity of transcription regulation (Kouzarides, 2007; Lawrence et al., 2016; Li et al., 2007). An appealing biological phenomenon is the co-occurrence of activating histone modifications and repressive histone modifications at the same chromatin region, namely bivalent chromatin (Bernstein et al., 2006). Bivalent chromatin is initially reported to mark development-related genes in embryonic stem cells, and further studies validate its existence in lineage-committed progenitor cells, tumor cells, and other cell types (Azuara et al., 2006; Bernstein et al., 2006; Chaffer et al., 2013; Kinkley et al., 2016; Matsumura et al., 2015). A prominent bivalent chromatin form is composed of both the activating modification H3K4me3 and the repressive modification H3K27me3 (trimethylation of lysine 4 or lysine 27 on histone H3), which are catalyzed by the Trithorax group (TrxG) proteins and Polycomb group (PcG) proteins respectively (Schuettengruber and Cavalli, 2009; Schuettengruber et al., 2011). Other forms of bivalent chromatin such as H3K4me3/H3K9me3 and H3K9me3/H3K36me3 are also observed (Matsumura et al., 2015; Mauser et al., 2017). Such bivalent chromatin is found to be associated with cellular phenotypical plasticity during cell differentiation and oncogenesis (Bernstein et al., 2006; Chaffer et al., 2013; Matsumura et al., 2015; Rugg-Gunn et al., 2010). For example, the bivalent chromatin is viewed as a poised state with bidirectional potential by maintaining key developmental genes at low expression levels and waiting for further activation or repression upon appropriate signals (Bernstein et al., 2006).

However, several key issues of the bivalent chromatin remain controversial. First, the pattern formation mechanism of bivalent chromatin is still unclear. Intuitively, bivalent chromatin results from the balance between TrxG proteins and PcG proteins (Voigt et al., 2013), as suggested by the observations that mutations or experimental perturbations tilting such balance dramatically reduce bivalent histone modifications (Béguelin et al., 2013; Shema et al., 2016). How the bivalent chromatin emerges from the interaction between TrxG and PcG still remains unknown. Second, a quantitative analysis of bivalent chromatin composition at the nucleosome level is lacking. Most of the bivalent chromatin domains are identified by using separate single-antibody ChIP experiments for each of the modifications. Consequently, it is not sufficient to depict the modification state of each nucleosome at the chromatin region, and it is unclear whether the two marks reside simultaneously on the same nucleosome. While genome-wide sequential ChIP or other similar techniques like co-ChIP and re-ChIP can detect bivalent nucleosomes carrying both two marks (Kinkley et al., 2016; Sen et al., 2016; Weiner et al., 2016), the extent to which the bivalent nucleosomes account for the bivalent chromatin remains to be investigated. Thirdly, the causal role of bivalent chromatin in mediating phenotypical plasticity remains largely controversial. Several experimental studies suggest that many bivalent chromatin domains are resolved upon differentiation and the expression levels of the corresponding genes change as well (Bernstein et al., 2006). While such coincidence establishes the association between the bivalent chromatin and its proposed function, the causality is difficult to confirm because perturbations of bivalent chromatin inevitably lead to the change of a broad range of genes, making it difficult to interpret the observations (Sneppen and Ringrose, 2019; Voigt et al., 2013).

Previous mathematical models of histone modifications mainly focus on the emergence of epigenetic bistability and spatial patterns as well as heritability (Dodd et al., 2007; Zhang et al., 2014). The theoretical works on modeling the bivalent chromatin state are still limited and oversimplified. For example, when modeling the H3K4me3/H3K27me3 bivalent chromatin, the intermediate, yet critical, modification steps such mono- and dimethylations are usually neglected (Huang and Lei, 2017; Ku et al., 2013; Sneppen and Ringrose, 2019), which have been implicated to dramatically affect the system behavior by increasing nonlinearity (Berry et al., 2017). A comprehensive model has been developed to incorporate four kinds of histone modifications; however, details like the mutual exclusiveness of H3K4me3 and H3K27me3 on the same histone tail as well as different nucleosome interaction ranges are ignored, which may be critial to the bivalent dynamics (Sneppen and Ringrose, 2019).

In this paper, we develop a minimal yet fully equipped mathematical model to imitate the stochastic dynamics of two antagonistic histone modifications H3K4me3 and H3K27me3. Our model incorporates the essential elements, including the intermediate steps of histone modifications, the mutual exclusiveness of H3K4me3 and H3K27me3 on the same histone tail, and different nucleosome interaction ranges. To decipher the formation mechanism of bivalency, we investigate the parameter constraints that yield the bivalent chromatin state and analyze the biological importance of those constraints. We demonstrate that advantageous methylating activity over demethylating activity, noisy conversions of modifications, and nonlinearity are required for the emergence of bivalent chromatin. To probe the relationship between bivalent nucleosome and bivalent chromatin, we analyze the composition of bivalent chromatin at the nucleosome level under different nucleosome interaction regimes and predict that no more than 50% of nucleosomes at the bivalent chromatin domain are bivalent nucleosomes. Finally, by analyzing the transitions among chromatin states in response to external stimulus or noise, we find that bivalent chromatin allows the transition to multiple states and can serve as a stepping stone to facilitate a step-wise transition between repressive chromatin state and activating chromatin state, showing the potential role of bivalent chromatin in mediating phenotypical plasticity during cell fate decision.

## Results

### Modeling H3K4me3/H3K27me3 bivalent chromatin

To capture the details of histone modification patterns, we propose a minimal model while incorporating all the essential interaction elements. Inspired by a previous work (Berry et al., 2017), which focuses on H3K27 methylations for modeling the histone modification dynamics, our model is constructed to study the interplay between two distinct kinds of histone modifications, i.e., H3K4 methylations and H3K27 methylations.

In our model, a linear array of nucleosomes as well as their interactions, representing a chromatin region in the cell, is considered (Figure 1A). While each nucleosome actually is an octamer composed of two copies of four core histones (H2A, H2B, H3 and H4), we focus on the modifications on H3 histone and assume that a nucleosome consists of two copies of H3 histones by neglecting the other three types of histones for simplification. Each histone H3 is assumed to have one modification site, and can be in one of the seven modification states: K27me3, K27me2, K27me1, Empty, K4me1, K4me2 and K4me3 (Figure 1B). Here “Empty” means no methylation on histone H3. “Implicit” in the above assumption is that H3K27 methylations and H3K4 methylations are mutually exclusive on the same histone tail, due to the experimental observation that H3K27 methylations and H3K4 methylations can hardly exist on the same histone tail (Shema et al., 2016; Voigt et al., 2012).

**Figure 1.**
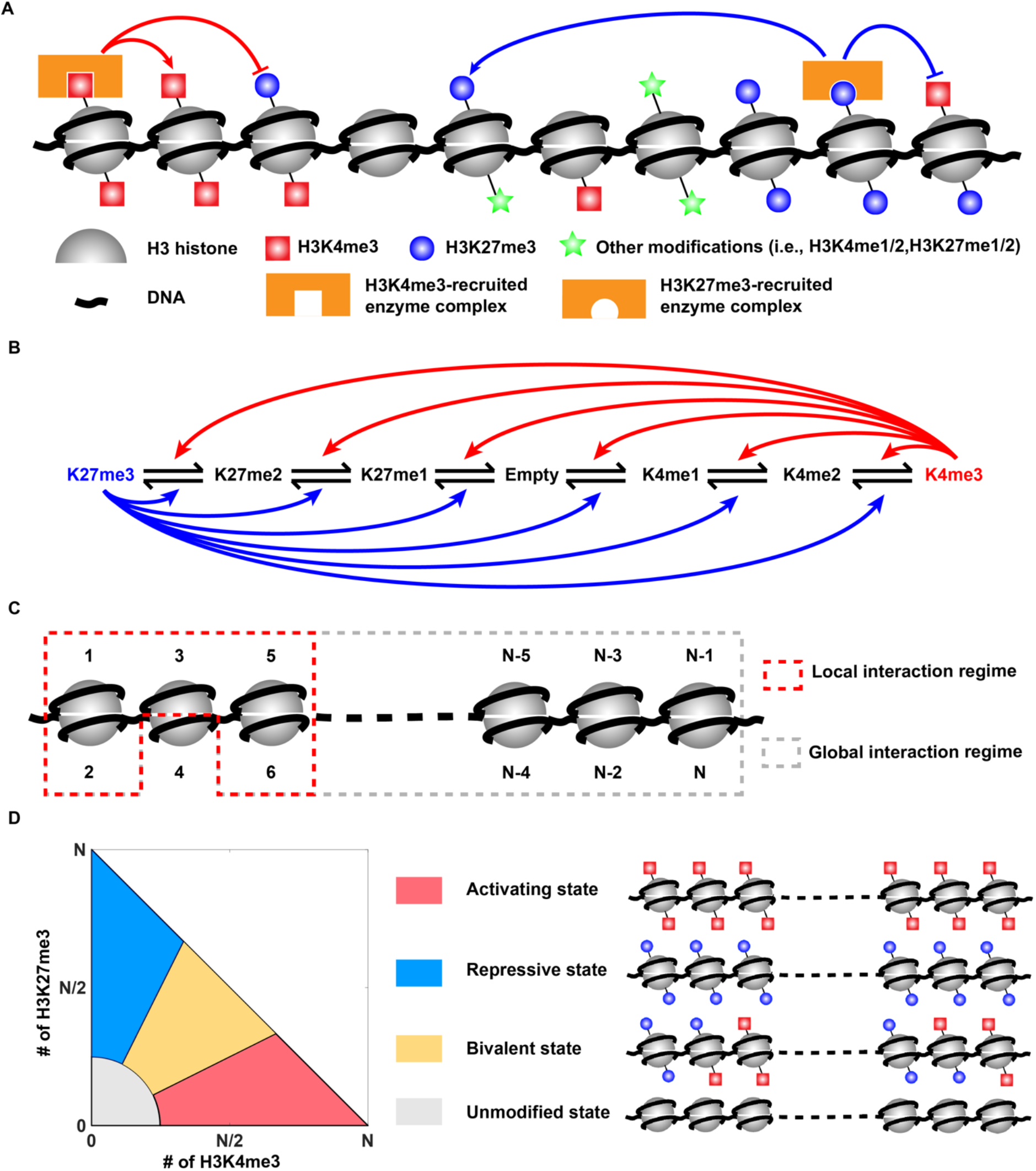
A model of histone modification at bivalent chromatin domain. (A) Schematic illustration of the model for histone modification dynamics. (B) The modification state of a histone H3 and feedbacks mediated by H3K4me3 and by H3K27me3. (C) An illustration of histone “neighbors” for local interaction regime and global interaction regime. For the local interaction regime, the neighbors of the 4th histone are histone number 1 2, 3, 5, and 6 (indicated by the red dashed box); for the global interaction regime, the neighbors include all the other *N*-1 histones (indicated by the gray dashed box). (D) Classification of the chromatin states. Four chromatin states are defined according to both the number of H3K4me3 and the number of H3K27me3. In the left plot, the region for the unmodified state is bounded by the curve *x^2^* + *y*^2^ = (*N*/4)^2^ and the axes in the first quadrant. The boundary between the regions for the bivalent state and the activating state is *y* = *x*/2. and the boundary between regions for the bivalent state and the repressive state is *y* = 2*x*. The right panel illustrates the possible composition for four chromatin states at the nucleosome level.

For each histone H3, the modification state can change towards neighboring states through one-step reactions (Figure 1B). While there are 12 possible reactions in total, at most two possible reactions can occur for a histone with a specific modification state. For example, if the current modification state of the *i*th histone (*i*=1, 2,…, *N,* where *N*is the total number of histones) is H3K4me2, then it can be either methylated to H3K4me3 or demethylated to H3K4me1. The reaction rate depends on not only the background catalytic activity of histone modification enzymes but also the modifications on neighboring nucleosomes. For instance, existing H3K27me3 can facilitate H3K27 methylations on neighboring nucleosomes by recruiting the enzymatic complex PRC2 (polycomb repressive complex 2) (Hansen et al., 2008; Margueron et al., 2009), and also enhance H3K4 demethylation by interacting with KDM5A through PRC2 (Pasini et al., 2008), thus forming a positive feedback for H3K27me3 deposition (Figure 1A). Similarly, a recruitment-mediated positive feedback for H3K4me3 deposition is also suggested by previous studies (Agger et al., 2007; Dou et al., 2006; Issaeva et al., 2007; Lee et al., 2007; Ruthenburg et al., 2007; Voigt et al., 2013; Wysocka et al., 2005). Thus, our model incorporates the feedbacks mediated by H3K4me3 and H3K27me3, by assuming that H3K4me3 (or H3K27me3) promotes all the six reactions towards H3K4me3 (or H3K27me3) on neighboring nucleosomes (Figure 1B). Besides, different nucleosome interaction ranges are considered when modeling the feedbacks from neighboring nucleosomes. In our model, by specifying the range of “neighbors” of each histone along with the assumption that only neighbors can interact with each other, we can establish different nucleosome interaction regimes. Two typical interaction regimes are studied: the local interaction regime where the neighbors of one histone include the other histone on the same nucleosome and histones in adjacent nucleosomes, and the global interaction regime where the neighbors include all the other histones regardless of their distance in the one-dimension chromatin fiber (Figure 1C).

The formulas used to model all 12 possible reactions for the *i*th histone are presented in Table 1. For the purpose of parameter reduction, the same parameters are used for the three (de)methylation reactions on H3K4 or H3K27 site. Each reaction rate contains a feedback term that reflects recruitment-mediated (de)methylation, and a noise term representing noisy (de)methylation resulting from background enzyme catalytic activity. For example, for the *i*th histone carrying H3K4me2, the methylation rate and demethylation rate can be formulated as *k*_K4me_*f*(*E_i_*) + *α*_K4me_ and *d*_K4me_*g*(*F*_i_) + *β*_K4me_, respectively. Here, *E_i_* and *F_i_* are defined as fractions of H3K4me3 and H3K27me3 among the neighbors of the *i*th histone, respectively. The methylation rate consists of a noise term *α*_K4me_ and a feedback term *k*_K4me_*f*(_*i*_), where *k*_K4me_ is the recruitment-mediated H3K4 methylation rate constant and *f*(*E_i_*) reflects the positive feedback mediated by neighboring H3K4me3 on the deposition of H3K4 methylations. Likewise, the demethylation rate contains a noise term *β*_K4me_, and a feedback term *d*_K4me_*g*(*F_i_*) representing the inhibitory effect of neighboring H3K27me3 on H3K4me3 by enhancing the removal of H3K4 methylations. Functions *f* and *g* are chosen as Hill functions to consider the nonlinear feedback effect. The rate formulas for all histones are constructed in a similar way (see Methods for more details).

**Table 1.** Rate formulas of all 12 reactions for the *i*th histone.

| Reactions | Reaction rate |
| --- | --- |
| Empty → H3K4me1, H3K4me1 → H3K4me2,<br>H3K4me → H3K4me3 | $k_{K4me}f(E_i) + \alpha_{K4me}$ |
| H3K4me3 → H3K4me2, H3K4me2 → H3K4me1,<br>H3K4me1 → Empty | $d_{K4me}g(F_i) + \beta_{K4me}$ |
| Empty → H3K27me1, H3K27me1 → H3K27me2,<br>H3K27me2 → H3K27me3 | $k_{K27me}g(F_i) + \alpha_{K27me}$ |
| H3K27me3 → H3K27me2, H3K27me2 →<br>H3K27me1, H3K27me1 → Empty | $d_{K27me}f(E_i) + \beta_{K27me}$ |

All parameters in our model are shown in Table S1. There are four free parameters in total, including *k*_K4me_, *k*_K27me_, *γ*_K4_ and γ_K27_. Since we fix recruitment-mediated demethylation rate constants *d*_K4me_ and *d*_K27me_ as ones, parameters *k*_K4me_ and *k*_K27me_ can be interpreted as the ratio of recruitment-mediated methylation rate and demethylation rate for H3K4 and H3K27, respectively, also called methylation-to-demethylation ratios. Parameters *γ*_K4_ and *γ*_K27_ are noise-to-feedback ratios, defined as the ratio of noisy (de)methylation rate to recruitment-mediated (de)methylation rate constant for H3K4 and H3K27, i.e., *γ*_K4_ = *α*_K4me_/*k*_K4me_ = β_K4me_/*d*_K4me_, γ_K27_ = *α*_K27me_/*k*_K27me_ = *β*_K27me_/*d*_K27me_, respectively.

### Emergence of the bivalent chromatin state

By direct simulating the system with Gillespie algorithm (Gillespie, 1977), we obtain the temporospatial dynamics of histone modifications. The chromatin states can be evaluated based on the abundance of H3K4me3 and H3K27me3 over the chromatin region. We classify the chromatin states into four categories, including activating chromatin state, repressive chromatin state, bivalent chromatin state and unmodified state (Figure 1D). If both H3K4me3 number and H3K27me3 number are at low levels, i.e., most of the histones are mono/dimethylated or unmodified, such chromatin state is named as the unmodified state. Otherwise, the chromatin is regarded as the activating state if the number of H3K4me3 is twice larger than that of H3K27me3, the repressive state if the number of H3K27me3 is twice larger than that of H3K4me3, or the bivalent state if the numbers of H3K4me3 and H3K4me27 are balanced (Figure 1D).

Under different parameter settings, our model can exhibit three distinct modes: monostability, bistability, and tristability. This means that the four chromatin states can either exist solely or coexist with others. For example, there exists only the repressive chromatin state when *k*_K27me_ is much greater than *k*_K4me_ (Figures 2A–2C), and the system can behave as bistability switching between the activating state and the repressive state when *k*_K27me_ is close to *k*_K4me_ (Figures 2D–2F). Besides, under proper parameter settings, the bivalent chromatin state can also be observed in a tristable mode, coexisting with the activating state and the repressive state. Simulation of the stochastic model and analysis of the corresponding deterministic model both confirm such tristability (Figures 2G–2I).

**Figure 2.**
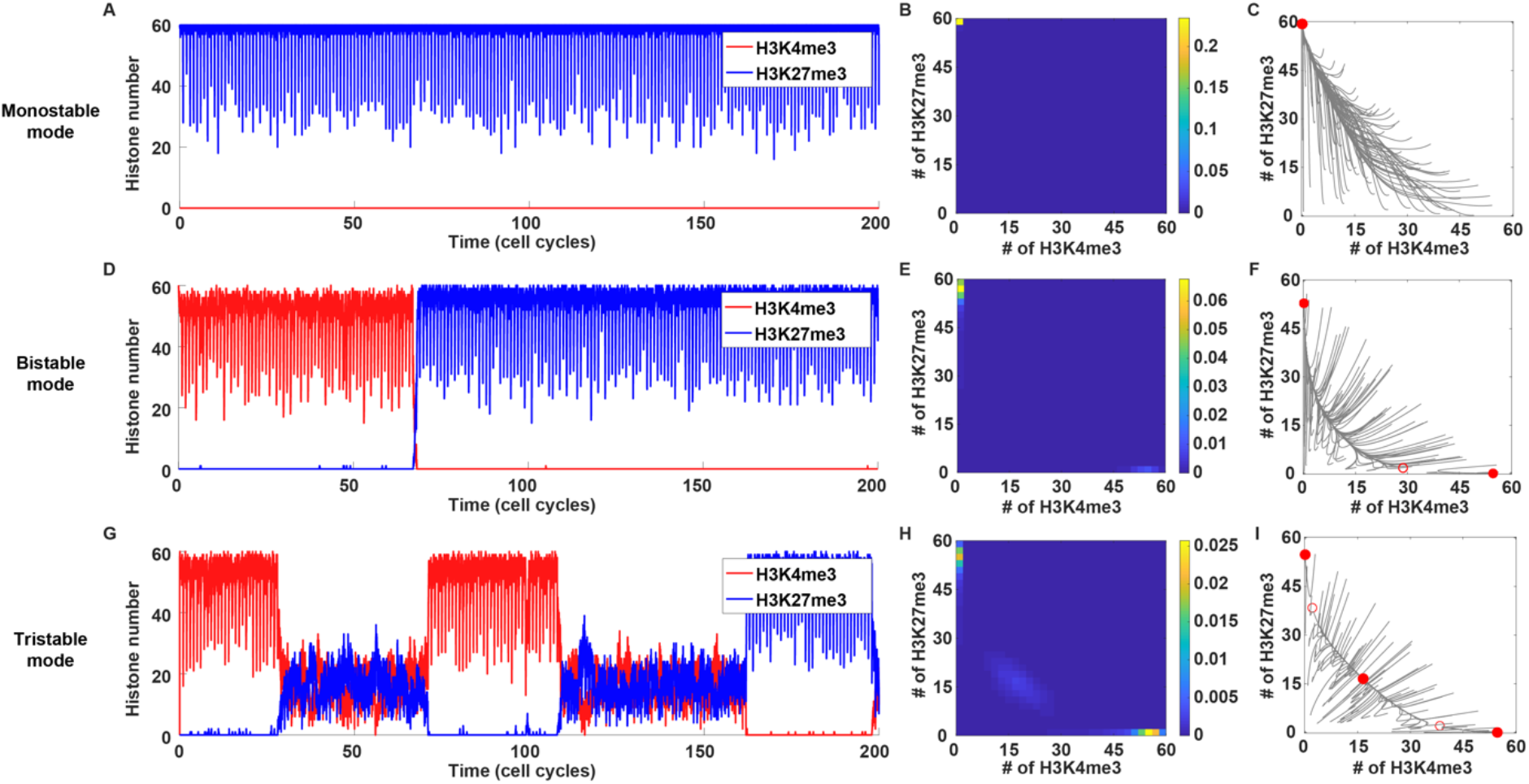
Typical simulations of histone modification dynamics and emergence of the bivalent chromatin state. (A-C) The dynamics (A), landscape (B) and phase plane (C) of a system behaving as the monostable repressive chromatin state. Parameters: *k*_K4me_ = 1, *k*_K27me_ = 10, *γ*_K4_ = *γ*_K27_ = 0.1. (D-F) The dynamics (D), landscape (E) and phase plane (F) of a system operating in a bistable mode where activating state and repressive state coexist. Parameters: *k*_K4me_ = 1, *k*_K27me_ = 1.6, *γ*_K4_ = *γ*_K27_ = 0.1. (G-I) The dynamics (G), landscape (H) and phase plane (I) of a tristable system where activating state, repressive state and bivalent state coexist. Parameters: *k*_K4me_=*k*_K27me_ = 2, *γ*_K4_ = *γ*_K27_ = 0.2. The two-dimensional chromatin state landscape (B, E, and H) is obtained by the density of stochastic trajectories, and the two-dimensional phase plane (C, F and I) is obtained by the deterministic trajectories (gray lines), where red solid dots and red circles represent stable states and saddles respectively. The global interaction regime is adopted in all stochastic simulations.

### Formation mechanisms of the bivalent chromatin state

#### 1) High methylation-to-demethylation ratio and high noise-to-feedback ratio

We next investigate the parameter space to answer what conditions are required for generating the bivalent chromatin state. First, we set the noise-to-feedback ratios for H3K4 and H3K27 are the same, i.e., *γ*_K4_ = *γ*_K27_ = *γ*. For a given level of noise-to-feedback ratio *γ*, we next calculate the time-averaged probabilities of four chromatin states under different combinations of *k*_K4me_ and *k*_K27me_ (Figures 3A and 3B). We find that, no matter which interaction regimes (local or global) are utilized, for most of the parameter combinations, the system exhibits monostable mode or bistable mode, a result consistent with previous studies (Berry et al., 2017; Dodd et al., 2007; Sneppen and Ringrose, 2019). Nevertheless, we still find that a relatively small parameter space exists for the bivalent state. When the methylation-to-demethylation ratios for H3K4 and H3K27 are well matched and both large enough, the bivalent state is usually observed in a tristable mode where the activating state and the repressive state as well as the bivalent state coexist. The requirement of high methylation-to-demethylation ratio (i.e., advantageous methylating activity over demethylating activity) for the bivalent state, is further confirmed by analyzing the deterministic version of our stochastic model (Figure S1). Compared with the global interaction regime, the local one shows a lower probability of the bivalent state but allows the bivalent state under smaller methylation-to-demethylation ratios (Figures 3A and 3B). In addition to the methylation-to-demethylation ratio, the noise-to-feedback ratio plays a role in the emergence of the bivalent state. High noise-to-feedback ratio, which reflects frequent noise conversions of histone modifications, leads to the large probability of the bivalent state; with the increase of noise-to-feedback ratio, the monostable bivalent state may emerge (Figures 3A and 3B). These results also hold for a small system size *N*=20 (Figure S2). Furthermore, more frequent cell division which could induce random loss of histone modifications also contributes to a higher probability of bivalent chromatin state (Figure S3).

**Figure 3.**
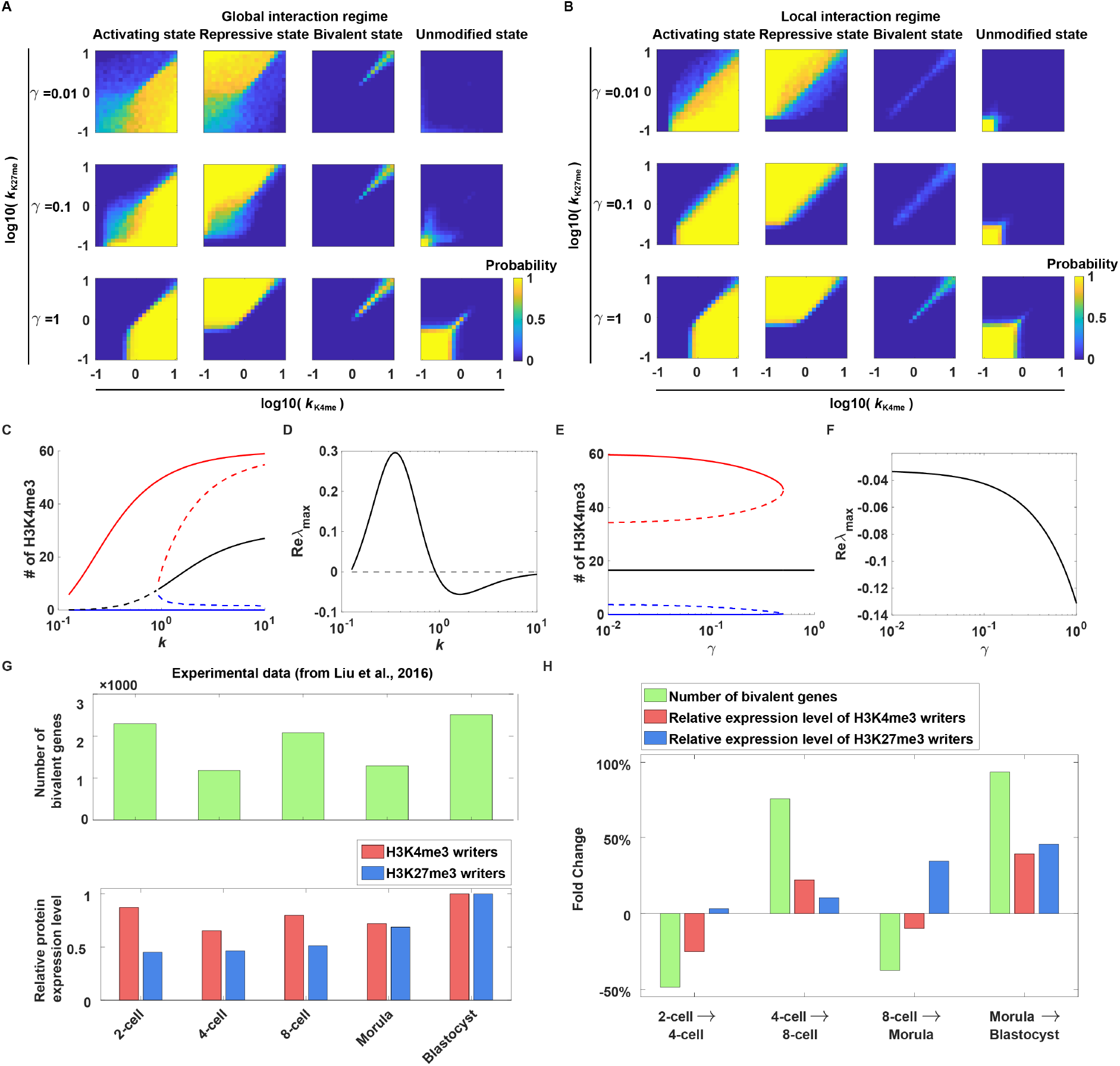
The effects of methylation-to-demethylation ratio and noise-to-feedback ratio on the emergence of bivalent chromatin state. (A-B) Parameter space for four different chromatin states under global interaction regime (A) and local interaction regime (B). For each interaction regime, time-averaged probabilities of four chromatin states under different combinations of *k*K4me and *k*K27me are calculated. For each parameter set, 100 simulations with randomly assigned initial values are used for averaging; for these initial values, numbers of H3K4me3 and H3K27me3 obey a discrete uniform distribution on the region *D* = {(*x,y*)|*x* + *y* ≤ *N,x,y* = 0,1,…,*N*} and the other histones are set to be Empty. (C) The bifurcation diagram for the steady-state H3K4me3 number as a function of the methylation-to-demethylation ratio. Solid lines and dashed lines represent stable steady states and unstable steady states, respectively. (D) The maximum of the eigenvalue real parts of the Jacobean matrix for the intermediate steady state (indicated by the black line in (C)). (E-F) The same plots as (C-D) except that noise-to-feedback ratio *γ* is set to be the control parameter. Parameters for (C-D) are set to be *γ* = *γ*_K4_ = *γ*_K27_ = 0.2, while those for (E-F) are set to be *k* = *k*_K4me_ = *k*_K27me_ = 2. (G) The number of bivalent genes (upper panel, extracted from Extended Data Figure 8b of a previous study (Liu et al., 2016)) and the relative protein expression level of H3K4me3 and H3K27me3 writers (lower panel, extracted from Extended Data Figure 2d of a previous study (Liu et al., 2016)) in different embryo stages. The number of bivalent genes for the blastocyst stage embryo is estimated by the average of the numbers for the inner cell mass and trophectoderm. The median values of the relative protein expression levels of H3K4me3 and H3K27me3 writers are used and then normalized to the max among different stages. (H) The fold change of the number of bivalent genes and the relative protein expression levels of H3K4me3 and H3K27me3 writers between adjacent stages.

By utilizing the corresponding deterministic version of our stochastic model, we analyze how the methylation-to-demethylation ratio and the noise-to-feedback ratio affect the bivalent chromatin state. As shown in the bifurcation diagram in Figure 3C, the bivalent state derives from a branch between the activating state branch and the repressive state branch, i.e., an intermediate state. When the methylation-to-demethylation ratio (*k* = *k*_K4me_ = *k*_K27me_) is small, such intermediate state behaves as an unmodified state since the number of H3K4me3 (and H3K27me3) is close to zero. As *k* increases, the bivalent state emerges due to the increased amounts of H3K4me3 and H3K27me3. We also notice the non-monotonic effect of the methylation-to-demethylation ratio on the stability of the bivalent state: the intermediate state becomes stable once *k* exceeds some threshold, but the stability will reach its maximum and then declines as *k* increases beyond (Figure 3D). With the methylation-to-demethylation ratio is fixed, increasing the value of noise-to-feedback ratio *γ* has no effect on the H3K4me3 (or H3K27me3) number of the bivalent state but can stabilize the bivalent state (Figures 3E and 3F). Thus, we conclude that both the high methylation-to-demethylation ratio and the high noise-to-feedback ratio collectively contribute to the emergence of the bivalent chromatin state: high methylation-to-demethylation ratio pushes the system away from the unmodified state but towards the bivalent state, while high noise-to-feedback ratio stabilizes the bivalent state.

A recent experimental study shows the positive role of a high methylation-to-demethylation ratio in generating bivalent chromatin state (Liu et al., 2016). Liu et al. performed a genomewide analysis of H3K4me3 and H3K27me3 in mouse pre-implantation embryos and observed that the number of bivalent genes change dynamically over different embryo developmental stages (the upper panel of Figure 3G). The relative protein expression levels of H3K4me3 and H3K27me3 writers are measured (the lower panel of Figure 3G). By analyzing how the number of bivalent genes and expression levels of H3K4me3 and H3K27me3 writers change over adjacent stages, we find that simultaneous elevation of expression levels of antagonistic modification writers is associated with the increased number of bivalent genes (Figure 3H). This is consistent with our finding that the higher methylation-to-demethylation ratios lead to the more probable bivalent chromatin state.

#### 2) Nonlinearity

As mentioned above, the bivalent chromatin state usually emerges along with the tristability, which typically results from the nonlinearity. The nonlinearity in our model arises not only from the Hill functions (vs linear functions) to describe the feedback, but also by incorporating multiple positive feedback loops in the reactions converting between H3K4me3 and H3K27me3. For example, the overall rate for consecutive conversions from H3K27me3 to H3K4me3 is proportional to (*d*_K27me_*f*(*E_i_* + β_K2727me_)^3^(*k*_K4me_*f*(*E_i_*) + *α*_K4me_)^3^, a nonlinear function naturally generated by the multiple positive feedback loops mediated by H3K4me3.

Here, we show that sufficient amount of nonlinearity is necessary to generate the chromatin tristability, leading to the bivalent state. To elucidate the effect of nonlinearity, we construct 11 variants of the standard model with weakened nonlinearity by reducing intermediate histone modification states, eliminating the feedback on either methylation reactions or demethylation reactions, or using the linear feedback, and then investigate their robustness of generating tristability (Figures 4A and 4B). For each of the 12 models (one standard model plus 11 variants), the robustness is measured by Q value (Qiao et al., 2019), defined as the number of parameter sets that yield chromatin tristability among 10,000 sets of randomized parameters by simulating the deterministic dynamics. Compared with the standard model, these variants show decreased Q values which mean reduced robustness of generating tristability (Figure 4C). Although some variants with the linear feedback can achieve tristability, the intermediate state of such tristability is usually not the bivalent state but the unmodified state (Figure S4). In addition, enhancing nonlinearity by increasing the Hill coefficient within a certain range dramatically improves the Q value for the standard model (Figure 4D). Since high Hill coefficients usually reflect strong cooperativity, a nonlinear phenomenon where the production rate of one species (e.g., H3K4me3) could become more dependent on the amount of itself as its amount increases, our result also indicates that high cooperativity is required for robust tristability. Taken together, these results illustrate the key role of nonlinearity in generating tristability.

**Figure 4.**
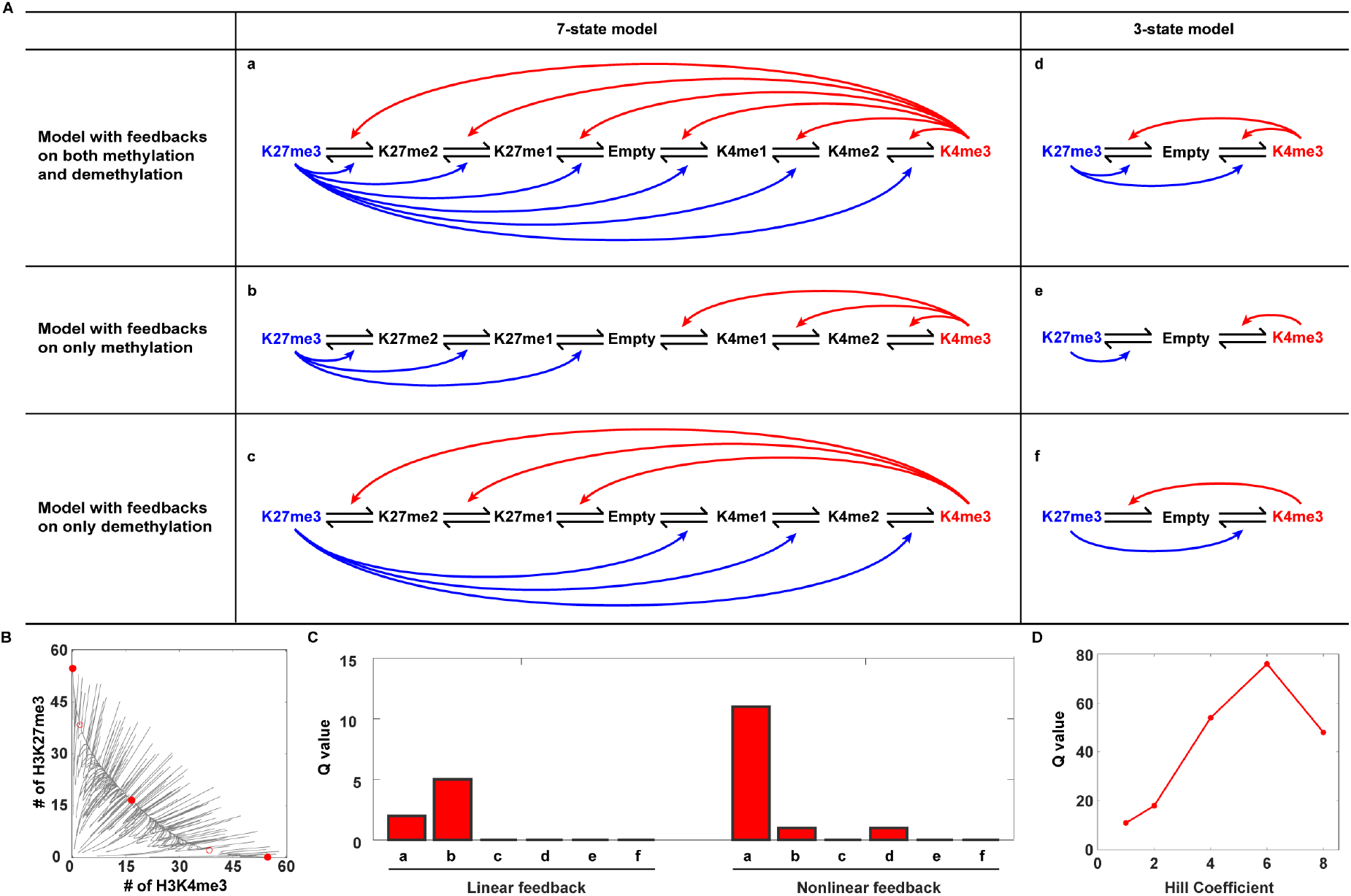
The role of nonlinearity in generating the bivalent state. (A) The standard model and its variants with reduced nonlinearity. These models include (a) 7-state model with feedbacks on both methylation and demethylation, (b) 7-state model with feedbacks on methylation only, (c) 7-state model with feedbacks on demethylation only, (d) 3-state model with feedbacks on both methylation and demethylation, (e) 3-state model with feedbacks on methylation only, and (f) 3-state model with feedbacks on demethylation only. Each of (a-f) can use either nonlinear feedback or linear feedback, thus forming 12 models in total. For the linear feedback, we choose *f(x)* = *g(x)* = *x*. (B) The typical phase plane of a system with tristability. (C) Q values of 12 models (including the standard model and 11 variants). For each model, we use the Latin hypercube method to sample parameters from loguniform distributions with the parameter ranges [0.1,10] for *k*_K4me_ and *k*_K27me_, [0.01 100] for *γ*_K4_ and *γ*_K27_, and [0.001,1000] for *K_f_* and *K_g_*. (D) The Q value as a function of the Hill coefficient for the standard model. Here, Hill functions *f* and *g* have the form of *f*(*x*) = *x^n^*/(*x^n^* + *K_f_^n^*) and *g*(*x*) = *x^n^*/(*x^n^* + *K_g_^n^*), respectively, where *n* is the Hill coefficient.

### Composition of bivalent chromatin at the nucleosome level

While the chromatin region exclusively occupied by bivalent nucleosomes which contain H3K4me3 and H3K27me3 on the opposing H3 histone tails is obviously bivalent, the inverse is not always true. To clarify the relationship between the bivalent chromatin and the bivalent nucleosome, we investigate the composition of the bivalent chromatin state at the nucleosome level. We find that the proportion of bivalent nucleosomes at the bivalent chromatin depends on what kind of interaction regime is applied (Figure 5). Under the global interaction regime, the bivalent state shows the pattern of randomly arranged H3K4me3 and H3K27me3 along the chromatin (Figure 5A), and the numbers of nucleosomes carrying H3K4me3/H3K4me3, H3K4me3/H3K27me3, H3K27me3/H3K27me3 at the bivalent chromatin are approximately in the ratio 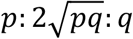 (Figures 5B and 5C). Furthermore, such ratio could be explained by a trinomial distribution with two parameters 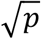 and 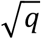 which represent probabilities for a histone to be H3K4me3 and H3K27me3, respectively (Figure S5). However, for the local interaction regime, the composition of the bivalent state is different: the chromatin region is composed of H3K4me3 patches and H3K27me3 patches (Figure 5D), and the ratio of the three kinds of nucleosomes becomes 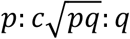, where 0 < *c* ≤ 2 (Figures 5E and 5F). This means that the proportion of bivalent nucleosomes shrinks under the local interaction regime compared with the case under the global interaction regime, which is because the local interaction regime promotes spatially gathering of the same histone modifications (i.e., H3K4me3 or H3K27me3). Nevertheless, the proportion of bivalent nucleosomes is further showed to be an increasing function of the noise-to-feedback ratio *γ*, and the upper bound 50% can be approached when extremely large *γ* is used (see the inset of Figure 5F).

**Figure 5.**
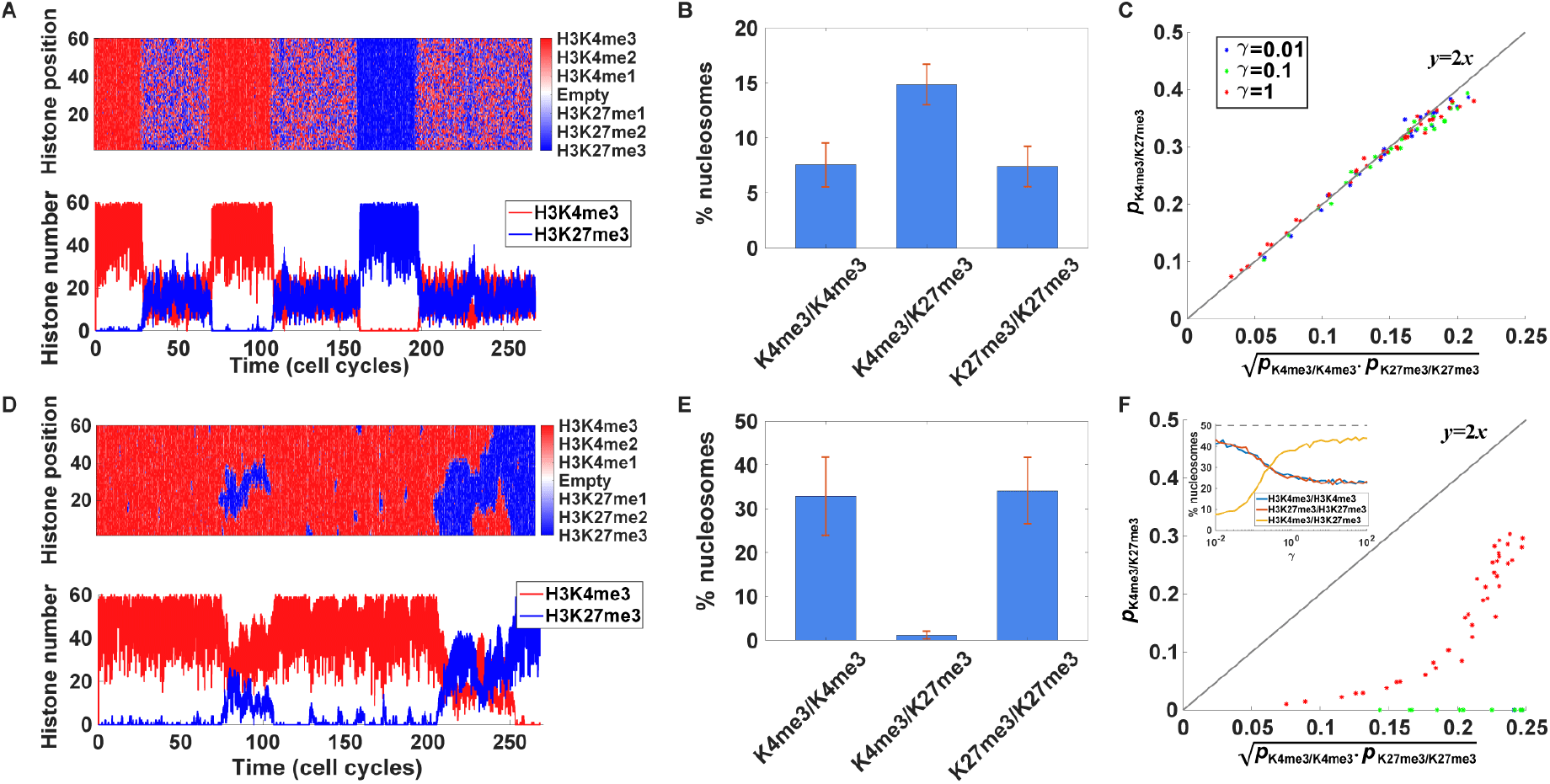
The composition of bivalent chromatin state at the nucleosome level under different interaction regimes. (A) The typical dynamics under global interaction regime. Upper panel: the temporospatial dynamics of histone modification. Lower panel: the dynamics of H3K4me3 number and H3K27me3 number. (B) The time-averaged percentages of different nucleosome types at the bivalent chromatin state. Only percentages of nucleosomes with H3K4me3/H3K4me3, H3K4me3/H3K27me3 and H3K27me3/H3K27me3 are shown. The data are shown as mean ± SD for 100 repeated and independent simulations. (C) The relationship between H3K4me3/H3K27me3 nucleosome percentage and (the square root of) the product of H3K4me3/H3K4me3 and H3K27me3/H3K27me3 nucleosome percentages. (D-F) The same plots as (A-C) except that local interaction regime is applied. Inset of (F): the percentage of bivalent nucleosomes at the bivalent chromatin under local interaction regime as a function of noise-to-feedback ratio. For (A-B) and (D-E), parameters are set to be *k*_K4me_ = *k*_K27me_ = 2 and *γ*_K4_ = *γ*_K27_ = 0.2. For (C) and (F), each data point represents a parameter set with randomly assigned *k*_K4me_ and *k*_K27me_ as well as a fixed *γ*, which yields the bivalent state. For the inset of (F), we set *k*_K4me_ = *k*_K27me_ = 20 and vary log_10_(*γ*) from −2 to 2 with an increment 0.1.

Taken together, an important finding is that the maximum proportion of bivalent nucleosomes at the bivalent chromatin is 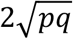 (or near 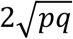 considering stochasticity due to the finite system size *N*) regardless of nucleosome interaction regimes. Moreover, 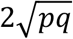 can reach its maximum value 50% when methylation-to-demethylation ratios are far greater than one and symmetric parameters are adopted. In particular, when methylation-to-demethylation ratios are far greater than one, histone modifications at bivalent chromatin are almost either H3K4me3 or H3K27me3, i.e., 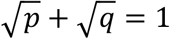; symmetric parameters further lead to *p* = *q* = 25%. Thus, 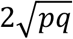 can reach a maximum of 50%.

It should be noted that, the fact that the proportion of bivalent nucleosomes cannot exceed 50% in our model results from mutual exclusiveness of K4 methylations and K27 methylations on the same H3 histone tail. Because of such mutual exclusiveness, a nucleosome can be in one of the only four modification states, i.e., H3K4me3/H3K4me3, H3K4me3/H3K27me3, H3K27me3/H3K27me3, and Empty/Empty, if we neglect the mono- and dimethylations. Then high methylation-to-demethylation ratio, required for the emergence of bivalent chromatin, will increase the proportions of all kinds of nucleosomes except Empty/Empty nucleosomes. Thus, H3K4me3/H3K27me3 nucleosomes must coexist with H3K4me3/H3K4me3 nucleosomes and H3K27me3/H3K27me3 nucleosomes. However, the case without such mutual exclusiveness could yield a bivalent state with the percentage of bivalent nucleosomes more than 50% and even near 100%. We remove the mutual exclusiveness in a modified model by assuming that each H3 histone has two separate modification sites, one for K4 methylations and the other for K27 methylations (Figures S6A and S6B). In this case, high methylation-to-demethylation ratio leads to a bivalent chromatin state, where near all H3K4 and H3K27 modification sites are occupied by H3K4me3 and H3K27me3 respectively; in other words, almost all nucleosomes are bivalent (Figure S6C).

### Bivalent chromatin as an intermediate state to mediate cell plasticity

Finally, we explore the functional roles of bivalent chromatin in mediating phenotypical plasticity during cell fate determination. To this end, we first investigate whether the bivalent chromatin state allows transitions to multiple states in response to either external stimulus or noise. To analyze how the bivalent state responds to the external stimulus, we perform a bifurcation analysis of the deterministic model, and find that the bivalent state can make transitions to other states as the control parameter (i.e., the external stimulus) changes (Figure 6A). When the methylation strength of H3K4 surpasses that of H3K27, the bivalent state switches to the activating state; when H3K4 methylation strength declines, it can switch to the repressive state. In the parameter regime of tristability without external stimulus, noise-driven transitions among the three chromatin states may spontaneously arise. For example, the noise caused by stochastic chemical reactions and the random loss of histone modifications during DNA replication, is found to be sufficient in triggering the transition from the bivalent state to either the activating state or the repressive state (Figure 5A). Furthermore, such plasticity of the bivalent state is found to be enhanced by cell division. As division occurs more frequently, the dwell time of the bivalent state becomes shorter, benefiting the switching to other states (Figure 6B). This result is significant because the division-enhanced plasticity only takes place for a subset of the intermediate states. For example, in the 7-state model with feedbacks on methylation only and with the linear feedback, there exists a tristable mode where the unmodified state serves as the intermediate state (Figure S4); however, the dwell time of the unmodified state increases as the cell division becomes faster (Figure 6C).

**Figure 6.**
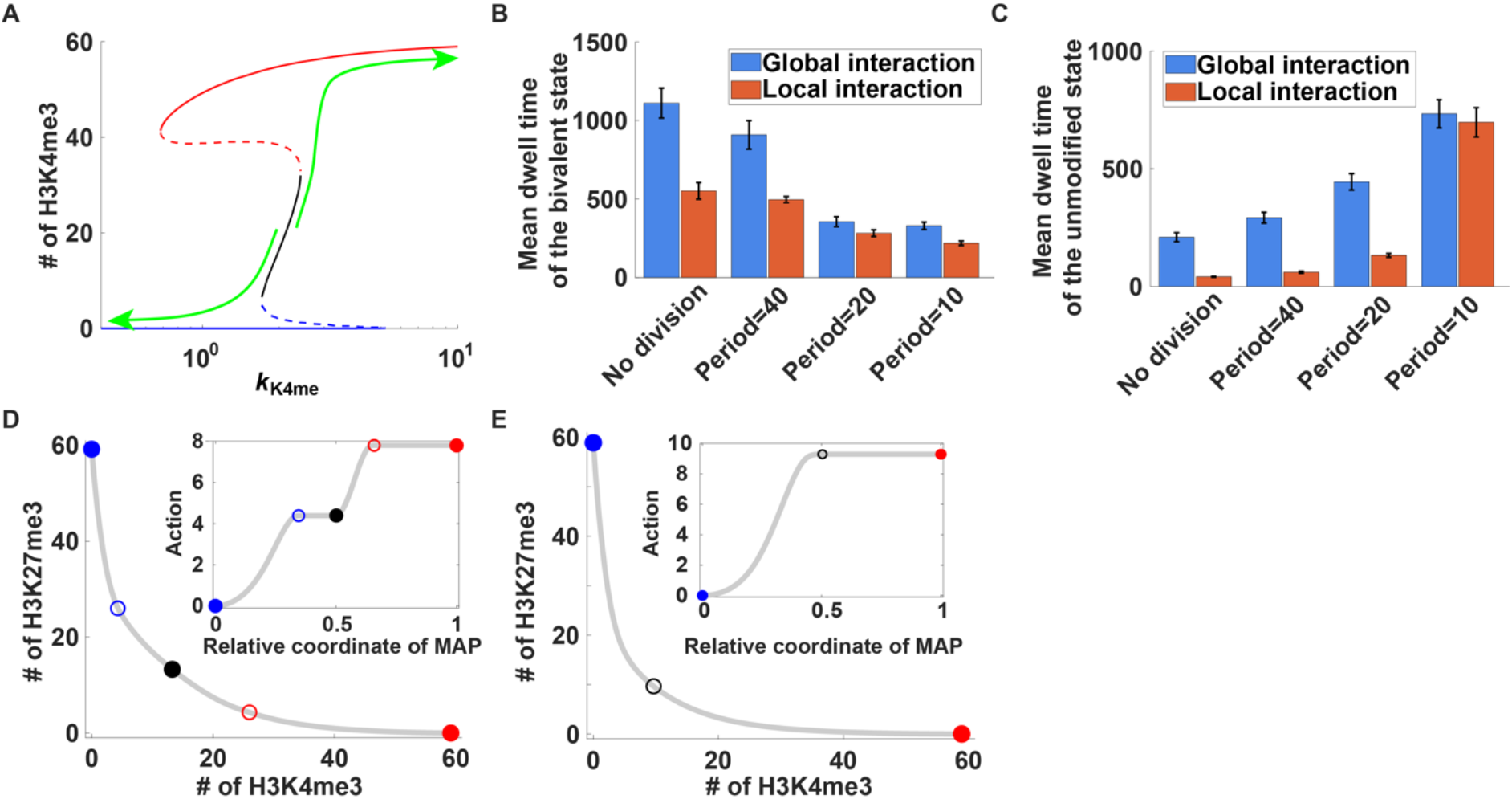
Bivalent chromatin as an intermediate state to mediate cell plasticity. (A) The bifurcation diagram of H3K4me3 number as a function of parameter *k*_K4me_. Parameters: *k*_K27me_ = 2 and *γ*_K4_ = *γ*_K27_ = 0.2. The green arrows illustrate the transition direction of the bivalent state as *k*_K4me_ changes. (B-C) The mean dwell time of the bivalent chromatin state (B) and the unmodified chromatin state (C) under the condition of no cell division, period=40, period=20, and period=10. The data are shown as mean±SD for multiple independent simulations, and the SD is obtained by using the bootstrap method. Parameters for (B) are set to be *k*_K4me_ = *k*_K27me_ = 2 and *γ*_K4_ = *γ*_K27_ = 0.1; for (C), the 7-state model with feedbacks on methylation only and linear feedback (see Figure 4Ab) is used, and parameters are set to be *k*_K4me_ = *k*_K27me_ = 5 and y_K4_ = y_K27_ = 0.04. (D-E) The MAP from the repressive chromatin state to the activating chromatin state for the parameter regime of tristability (D) and bistability (E). Inset: the accumulated action from the repressive chromatin state to the activating chromatin state along the relative coordinate ([0,1]) of the MAP. The MAP is calculated by using the geometric minimum action method (Heymann and Vanden-Eijnden, 2008), and 101 points are used in the discretization of the path. The dots in red, blue and black represent activating, repressive and bivalent state respectively; circles represent unstable steady states. Parameters: *γ*_K4_ = *γ*_K27_ = 0.02, *k*_K4me_ = *k*_K27me_ = 1.5 for (D) and *k*_K4me_ = *k*_K27me_ = 1.1 for (E).

In addition to the multi-directional transition potential, the bivalent chromatin can serve as an intermediate state, facilitating the noise-driven transitions between the repressive state and the activating state. We use the minimum action method (E et al., 2004; Heymann and Vanden-Eijnden, 2008; Yu et al., 2018) to compute the most probable path, i.e., minimum action path (MAP), for the transition. Interestingly, the MAP from the repressive state to the activating state passes through the bivalent state, indicating that the transition is a two-step process (Figure 6D). Compared to the repressive state, the bivalent state can facilitate the transition to the activating state because less action is required (see the inset of Figure 6D). Moreover, by comparing the entire action of the MAP for the cases with and without the bivalent state (i.e., tristability versus bistability), we find that the existence of the bivalent state can decrease the least action required for the transition between the two extreme chromatin states (Figures 6D and 6E). These results indicate that the bivalent state serves as a stepping stone to facilitate the step-wise transition between the two extreme chromatin states.

## Discussion

Bivalent chromatin has been associated with many important biological processes including cell differentiation and oncogenesis. However, the underlying mechanisms of the bivalent chromatin remain unclear. For instance, how does the bivalent chromatin emerge from the interaction between PcG proteins and TrxG proteins? To what extent do the bivalent nucleosomes account for the bivalent chromatin? In addition, could bivalent chromatin have a causal role in mediating phenotypical plasticity?

To address these questions, we developed a minimal but fully equipped model to perform a quantitative study of two antagonistic histone modifications H3K4me3 and H3K27me3. We identified that the high methylation-to-demethylation ratio and high noise-to-feedback ratio collectively contribute to the emergence of bivalent chromatin: high methylation-to-demethylation ratio pushes the system away from the unmodified state but towards the bivalent state, while high noise-to-feedback ratio stabilizes the bivalent state. Our finding is supported by a recent experimental observation that the increased number of bivalent genes is associated with simultaneous elevation of the expression levels of H3K4me3 and H3K27me3 writers (Liu et al., 2016). Dodd et al. have noticed that high noise-to-feedback ratio can increase the probability of intermediate states when studying the requirements for chromatin bistability (Dodd et al., 2007), which is consistent with the enhanced bivalent state stability induced by the high noise-to-feedback ratio. Besides, sufficient nonlinearity involved in our model, which has proved to be important for generating bistability (Berry et al., 2017; Dodd et al., 2007), could also extend the parameter space for generating multi-stability containing the bivalent state.

We quantitatively studied the composition of bivalent chromatin at the nucleosome level under different nucleosome interaction regimes: under the global interaction regime, proportions of different types of nucleosomes obey a trinomial distribution; under the local interaction regime, the proportion of bivalent nucleosomes shrinks, and is an increasing function of noise-to-feedback ratio. Most importantly, we predicted that bivalent nucleosomes account for no more than 50% of nucleosomes at the bivalent chromatin domain regardless of nucleosome interaction regime, which results from mutual exclusiveness of K4 methylations and K27 methylations on the same H3 histone tail. This result agrees with the scattered distribution of H3K4me3/H3K27me3 bivalent nucleosomes around the transcription start site of some bivalent genes (Sen et al., 2016). This could also explain why some of the previous models are difficult to produce a chromatin state with dominant bivalent nucleosomes (Huang and Lei, 2017; Ku et al., 2013; Sneppen and Ringrose, 2019). However, for the model without such mutual exclusiveness, bivalent nucleosomes can be dominant under proper parameters, which is consistent with a recent study (Sneppen and Ringrose, 2019). While removing the mutual exclusiveness is unrealistic for modeling H3K4me3/H3K27me3 bivalent chromatin, it could be feasible for other forms of bivalent chromatin.

Finally, by analyzing the transitions among chromatin states in response to external stimulus or noise, we demonstrated that bivalent chromatin can serve as a poised state, which has the potential to switch to multiple states and facilitates a step-wise transition between repressive chromatin state and activating chromatin state, to mediate phenotypical plasticity. Our results demonstrate why bivalent chromatin is often involved in many developmental processes where cell fate decision is required (Bernstein et al., 2006; Matsumura et al., 2015; Rugg-Gunn et al., 2010; Sachs et al., 2013). Such plasticity of bivalent chromatin also agrees with an experimental study wherein the bivalent gene Zeb1 can readily respond to microenvironmental signals and thus enables the transition to cancer stem cells (CSCs) from non-CSCs (Chaffer et al., 2013). However, the bivalent state is unlikely to be the only form of the poised chromatin. Several studies have suggested that a heterogeneous state such as bistability is the nature of the poised chromatin (Hong et al., 2011; Lorzadeh et al., 2016; Sneppen and Ringrose, 2019). This is not contradictory to our results. In fact, both the bivalent state and bistability reflect a balance between TrxG proteins and PcG proteins; from a mathematical point of view, under the condition with symmetric parameters, both the bivalent state and bistability can emerge, largely depending on the model and parameter setting. In addition, the bivalent state itself could contribute to a heterogeneous state because of its potential to mediate cell plasticity. Taken together, the combination of the bivalent state and the heterogeneous state could provide a full picture of the nature of the poised chromatin.

In the present model, we do not take into account the transcription processes, which have been implicated in the formation of bivalent chromatin (Voigt et al., 2013). Recent theoretical analyses have shown that incorporating the feedback from transcriptional products into the histone modification regulation can induce intermediate states (Huang and Lei, 2019; Zhang et al., 2019). So, it will be interesting to investigate the effect of transcription processes on the formation of bivalent chromatin in a model coupling both transcription processes and histone modification regulation. In addition, we expect that some new technology could be developed and experimentally test the predictions of our model. Although our model only focuses on H3K4me3/H3K27me3 bivalent chromatin, we expect that the underlying mechanisms are general and could be naturally applied to other combinations of histone modifications.

## Methods

### Stochastic Model

We consider a linear array of nucleosomes and each nucleosome has two copies of H3 histones. There are in total *N*histones, indexed by 1 to *N*. We also assume the (2*k-*1)th and *2k*th histones belong to the same nucleosome, where *k=1,*…, *N*/2. Let *S*_*i*_ denotes the modification state of the *i*th histone, and *S*_*i*_ ∈ {K27me3, K27me2, K27me1, Empty, K4me1, K4me2, K4me3}. In practice, we use seven numbers −3, −2, −1, 0, 1, 2, 3 to represent K27me3, K27me2, K27me1, Empty, K4me1, K4me2, K4me3 respectively.

The reaction rate, also termed as the propensity function, has been listed in Table 1. Despite 12 possible reactions in total, each histone *i* with state *S*_*i*_ can take part in at most two reactions, i.e., *S*_*i*_ → *S*_*i*_ + 1 and *S*_*i*_ →*S*, − 1. The propensity function for *S*_*i*_ → *S*_*i*_ + 1 is formulated as

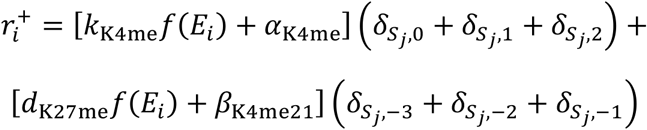

and the one for *S*_*i*_ → *S*_*i*_ − 1 is

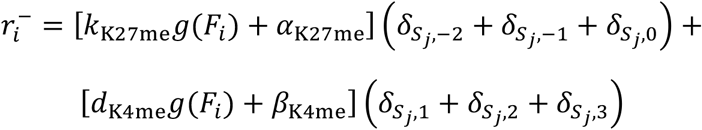

where 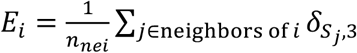 and 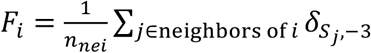 represent fractions of K4me3 and K27me3 among the neighbors of the *i*th histone respectively; *n*_nei_ is the number of histone neighbors; the function 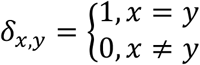 is the Kronecker delta. Functions *f* and *g* are chosen as Hill functions *f*(*x*) = *x*(*x* + *K_f_*) and *g(x)* = *x*/(*x* + *K_g_*) when modeling the nonlinear feedback, and chosen as *f*(*x*) = *g*(*x*) = *x* when modeling the linear feedback. The range of histone neighbors is defined according to the nucleosome interaction regime. For the global interaction regime, the neighbors are all the other *N*-1 histones, and *n*_*nei*_ = *N* − 1. For the local interaction regime, except four histones (number 1, 2 and *N*-1 and *N*) on the boundaries, there are 5 neighbors for each histone, i.e.,

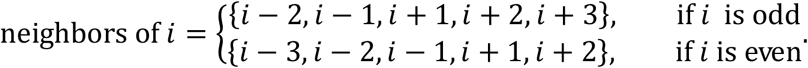

We set *n_nei_* = 5 for all histones for the local interaction regime.

We use Gillespie algorithm (Gillespie, 1977) to simulate the stochastic dynamics of the whole system. There are *N* species (i.e., histones) and 2*N* reaction channels in total. Denote *S*_*i*_(*t*)(*i* = 1,2,…,N) the state of each histone at time *t*, and denote vector ***S***(*t*) = (*S*_1_(*t*),*S*_2_(*t*),…,*S*_*N*_(*t*)) the state of the whole system. We denote *a*_2i-1_(*t*) and *a*_2*i*_(t) (*i* = 1,2,…,*N*) represent the two propensity functions for the *i*th histone: one for *S*_*i*_(*t*) → *S*_*i*_(*t*) + 1 and one for *S*_*i*_(*t*) → *S*_*i*_(*t*) − 1 respectively. The state change of the whole system induced by the *j*th reaction channel can be represent by a vector (***v***_*j*_ = (*j* = 1,2,…,2*N*) with the same length as ***S***(*t*), where the *j*th element is 1 if*j* is odd and −1 otherwise, and all the other elements are zeros. The Gillespie algorithm is as follows:

1. Initialization. Set the time variable *t* = 0, and assign desired values to ***S***(0).
2. Calculate and store the values of 2*N* propensity functions *a_j_*(*t*)(*j* = 1,2,…,2N) at current time *t*. Also calculate the sum of the 2*N* values, denoted as *a*_0_.
3. Generate two independent random numbers *r*_1_, and *r*_2_, both of which obeys a uniform distribution over the interval [0,1], and calculate the waiting time 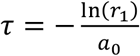 for the next reaction to occur. Determine the integer *μ* according to 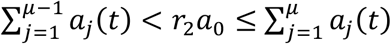. Here, *μ* is the index of reaction channel chosen to occur.
4. Update the time variable as *t* + *τ*, and update the system state as ***S***(*t* + *τ*) = ***S***(*t*) + *V_μ_*
5. Repeat the step 2)-4) until the terminating condition are satisfied.

Besides, DNA replication during the cell cycle is also simulated in our model. Denote *L* the cell cycle length. If the current time plus the waiting time for the next reaction exceeded the time when DNA replication should have occurred, i.e., *t* < *nL* ≤ *t* + *τ* for some positive interger *n*, then we stop the Gillespie algorithm and execute DNA replication. In particular, time variable *t* is updated as *nL* rather than *t* + τ, and each nucleosome will be replaced by an unmodified nucleosome (i.e., both *S*_2*k*-1_ (*nL*) and *S*_2*k*_(*nL*) are set to be zeros, *k*=1,…, *N*/2) with a probability of 0.5. After such DNA replication, resume the Gillespie algorithm by repeating the step 2)-4). We set the maximum number of iterations as 10^4^ for all stochastic simulations in our study unless otherwise specified.

### Deterministic model

Under the assumption of the homogeneous system and global interaction regime, the mean behavior of the stochastic model can be approximated by a set of ODEs. Denote *n*_K27me3_, *n*_K27me2_, *n*_K27me1_, *n*_Empty_, *n*_K4me1_, *n*_K4me2_ and *n*_K4me3_ the number of K27me3, K27me2, K27me1, Empty, H3K4me1, K4me2 and K4me3, respectively. Then the deterministic model can be described by ODEs as follows:

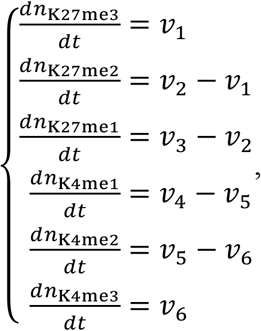

where

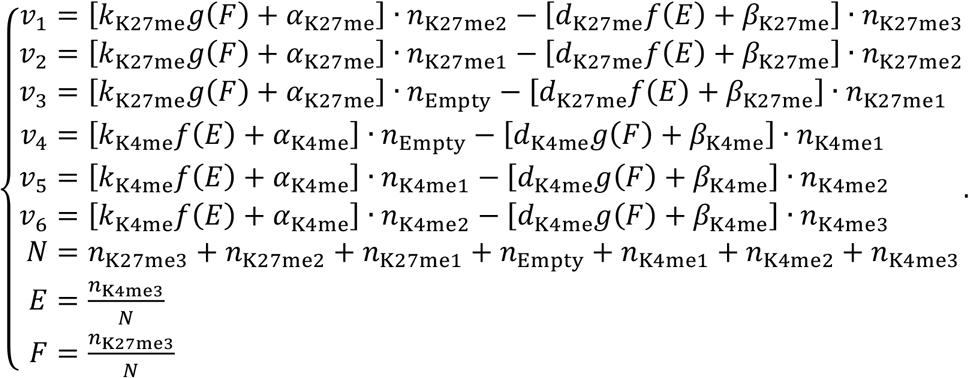

Similar ODEs can be constructed for the model variants shown in Figure 4.

## Acknowledgments

This work was supported by the National Natural Science Foundation of China No. 12050002. Q. N. is partly supported by a NSF grant DMS1763272, and The Simons Foundation grant (594598,QN).

## Author Contribution

L.Z. and Q.N. conceived the project; W.Z., L.Q. and S.Y. performed the analysis and computational simulations; L.Z. and Q.N. supervised the project; W.Z., L.Q., Q.N. and L.Z. wrote the paper.

